# The sociobiome–oral microbiome pathway in dental caries among Indigenous Australians

**DOI:** 10.1101/2025.09.16.676503

**Authors:** Sonia Nath, Laura S. Weyrich, Gina Guzzo, Joanne Hedges, Manisha Tamrakar, Kostas Kapellas, Lisa Jamieson

**Affiliations:** Adelaide Dental School, The University of Adelaide, Adelaide, SA, 5000, Australia; School of Biological Sciences, University of Adelaide, Adelaide, SA, 5000, Australia

**Author notes:** Current Address: Department of Anthropology and Huck Institutes of the Life Sciences, The Pennsylvania State University, University Park, PA, 16802, USA. Corresponding Author: Sonia Nath, Australian Research Centre for Population Oral Health Adelaide Dental School University of Adelaide, Adelaide, South Australia – 5000 Australia.

**Keywords:** oral microbiome, health inequities, socioeconomic status, dental caries, Aboriginal health

## Abstract

Indigenous Australians experience disproportionately high rates of dental caries, yet the biological pathways linking socioeconomic disadvantage to oral health remain unclear. This study examined how individual– and neighbourhood-level socioeconomic status (SES) shape the oral microbiome and mediate dental caries risk in Indigenous adults. A cross-sectional study of 100 Indigenous Australians (≥18 years) collected demographic, SES, and oral health behaviour data, followed by dental examinations for dental caries assessment, followed by collection of saliva and plaque samples. The microbiome was profiled using 16S rRNA sequencing, with analyses of microbial diversity, composition, differential abundance, and mediation of SES–caries associations. Saliva exhibited greater observed and Shannon diversity than plaque (both p < 0.01), with significant compositional differences (adonis p < 0.001). In saliva, alpha diversity was reduced with age, secondary education, low income, ownership of a healthcare card, and caries presence (all p < 0.05). SES explained greater variation in saliva than plaque composition, with associations for income (R²=3.8%, p<0.01), education (R²=2.0%, p<0.01), and caries (R²=2.2%, p<0.01). Differentially abundant taxa in low-income and caries groups included *Rikenellaceae RC9 gut group*, *F0058*, *Filifactor*, and *Treponema*. Mediation analyses showed 75.6% of the income effect on caries was mediated by microbiome shifts (ME=0.28, SE=0.32), compared with 21% for education (ME=0.03, SE=0.02). Socioeconomic disadvantage has a significant impact on the oral microbiome, influencing caries risk through income-related microbial dysbiosis. Saliva emerges as a sensitive biomarker of SES gradients. Addressing oral health inequities requires both structural policies targeting income inequality and microbiome-informed interventions.

**IMPORTANCE:** This research provides novel biological insights into how socioeconomic disadvantage contributes to the higher burden of dental caries among Indigenous Australians. Although social determinants of health are well recognised, the pathways connecting these determinants to oral disease remain unclear. By demonstrating that low income and education affect oral microbiome diversity and composition, and that a significant part of income-related caries risk is mediated through microbiome changes, this study highlights an important mechanism behind oral health inequalities. Identifying saliva as particularly responsive to socioeconomic differences makes it a useful, non-invasive biomarker for tracking risk in vulnerable populations. These findings emphasise the need for two approaches: structural interventions to reduce social and income gaps, and microbiome-based strategies to address microbial imbalance and disease risk. Together, they strengthen the evidence for more effective, culturally sensitive efforts to promote oral health equity.

## INTRODUCTION

Oral health disparities among Aboriginal and/or Torres Strait Islander Peoples (hereafter respectfully referred to as Indigenous people) represent a profound intersection of historical injustice, structural inequity, and biological vulnerability (1). Indigenous Australians experience disproportionately high rates of dental caries, periodontal disease, and tooth loss compared to non-Indigenous Australian populations (2, 3). These inequities are not a reflection of individual behaviour but are deeply rooted in the enduring legacies of colonisation and systemic racism that perpetuate socioeconomic disadvantage and restrict access to culturally safe oral healthcare (1). Neoliberal policies further entrench disparities by privatising dental care, reducing public health funding, and individualising responsibility for health outcomes, disproportionately marginalising Indigenous communities (4). This disparity cannot be explained by individual factors alone but must be understood through the lens of social determinants of health (SDOH) that shape oral health inequities.

While the biological mechanisms underlying caries development are well-documented, the role of SDOH in shaping the oral microbiome, a key mediator of caries, remains underexplored, especially in Indigenous Australian communities. The emerging concept of the “sociobiome”, the relationship between social determinants and microbial ecosystems, provides a powerful framework for understanding Indigenous oral health disparities (5, 6). According to the Australian Institute of Health and Welfare (AIHW), a framework for determinants of health conceptualises the complex causal pathway spanning upstream structural determinants, midstream behavioural factors, and downstream biological mechanisms (7). Socioeconomic status (SES) (8) is characterised by income, education, and occupation, which influence material circumstances such as housing conditions, neighbourhood characteristics, and financial capacity to access dental services (7, 9). Lower educational attainment and income are linked to elevated abundances of caries-associated taxa, such as *Lactobacillus* and *Prevotella*, alongside reduced microbial diversity, reflecting disparities in health literacy, dietary quality, and stress exposure (5, 10). Even with healthcare card ownership (indicative of receiving social welfare income and eligibility for subsidised services), access to preventive dental care may remain limited due to systemic barriers, perpetuating cycles of dysbiosis-driven conditions such as dental caries (11). Neighbourhood-level factors such as living in a remote location or a lower socioeconomic suburb may amplify these effects (5). Residing in socioeconomically deprived areas is associated with distinct oral microbial profiles, characterised by reduced evenness and overrepresentation of pathogenic genera such as *Porphyromonas* and *Fusobacterium* (5, 12).

For Indigenous Australians, these microbial disruptions are compounded by structural inequities: limited access to culturally safe dental care and intergenerational trauma from colonial policies (13, 14). Researchers found that Indigenous Australians had significantly higher oral microbial diversity and distinct composition than non-Indigenous individuals, including unique taxa not previously observed in human oral microbiota (14, 15). Similarly, work in Venezuela, the Philippines, and Uganda has reported systematic differences between the oral microbiota of Indigenous groups with traditional lifestyles and counterparts living agricultural or industrialised lifestyles (13). Despite growing recognition of these dynamics, critical gaps persist. The extent to which SES influence these microbial shifts remains poorly understood. Additionally, limited research has compared how saliva and plaque samples differentially capture these influences (16). To our knowledge, the role of the oral microbiome in relation to SES and dental caries has not been evaluated in Indigenous Australian adults. Therefore, this study aims to investigate how individual (e.g., education level, income source, ownership of healthcare card, dental affordability) and neighbourhood-level socioeconomic factors shape the composition and diversity of the oral microbiome in Indigenous populations.

## RESULTS

### Study population characteristics (Table S1)

The mean age was 41.92 years (SD=13.37), with mostly female participants (63.37%) and 91.0% identified as Indigenous. Age distribution showed that nearly half of the participants were in the 35-54 years category (47.0%), followed by 18-34 years (34.0%) and >55 years (19.0%). Data collection occurred almost equally across settings, with 55.6% at home/community centres. 58.0% of participants had secondary education or less, while 42.0% had tertiary education. Regarding income sources, 61.0% reported employment as their primary source, while 39.0% received Centrelink payments from the government. Healthcare card possession was evenly distributed (49.0% had cards, 51.0% did not). Financial hardship was prevalent, with 76.0% of participants reporting difficulty paying $100 for dental treatment. Most participants resided in metropolitan areas (86.0%) versus remote/regional locations (14.0%). According to the SEIFA Index, 59.0% lived in low socioeconomic status areas, while 41.0% lived in high SES areas. Dental care access was limited, with 66.0% reporting their last dental visit was over a year ago. Most (79.8%) visited dentists for problems rather than preventive check-ups (20.2%). Regarding health behaviours, 21% self-reported diabetes, and smoking status was distributed among current smokers (36.0%), non-smokers (38.0%), and past smokers (26.0%). These findings highlight significant socioeconomic disadvantage and barriers to dental care access within this population.

### Oral sample type influences oral microbiome diversity and composition

The analysis revealed significant microbial diversity and community composition differences between plaque and saliva samples. Observed features and Shannon’s diversity were significantly higher in saliva samples than plaque samples (p < 0.01) (Figure 1A). Beta diversity analysis clustering (Figure 1B) further demonstrated that microbial communities distinctly segregate by sample type (adonis p < 0.001), reflecting the unique ecological conditions and functional roles associated with plaque and saliva within the oral cavity (17, 18). A Venn diagram (Figure 1C) comparing core microbiome genera between plaque and saliva further highlighted these differences. Saliva contained 10 unique core genera, including *Megasphaera*, *Fretibacterium*, *Mogibacterium*, and *Stomatobaculum* and plaque samples had four unique core genera, which included *F0332*, *Kingella*, *Cardiobacterium* and *Aggregatibacter* and 22 core genera were shared between both sample types, including prominent taxa like *Campylobacter*, *Leptotrichia*, *Fusobacterium*, *Treponema*, *Prevotella*, and *Corynebacterium*. These shared genera represent key members of the oral microbiome that thrive across different niches but may exhibit functional specialisation depending on the environment. The microbial community composition bar plot revealed clear differences between the sample types (Figure 1D). Plaque samples were dominated by genera such as *Leptotrichia*, *Corynebacterium*, *Actinomyces*, *Fusobacterium,* and *Capnocytophaga*, which are characteristic of the localised biofilm environment of dental plaque. In contrast, saliva samples showed higher relative abundances of *Palleniella*, *Veillonella*, *Pasteurellaceae* family, and *Streptococcus*. Differential abundance analysis (Figure 1E) further emphasised the distinct microbial profiles with plaque associated with *Corynebacterium*, *F0332, Eikenella*, *Cardiobacterium*, and *Centipeda* and saliva associated with genera such as *Megasphaera*, *Solobacterium*, *Oribacterium*, *Palleniella*, and *Peptostreptococcus.* Plaque supports biofilm-associated microbes that are adapted to a localised anaerobic environment, while saliva has a broader range of microbial taxa influenced by its interactions with external factors and other oral microbial ecologies, such as the surrounding soft tissue.

**Figure 1.**
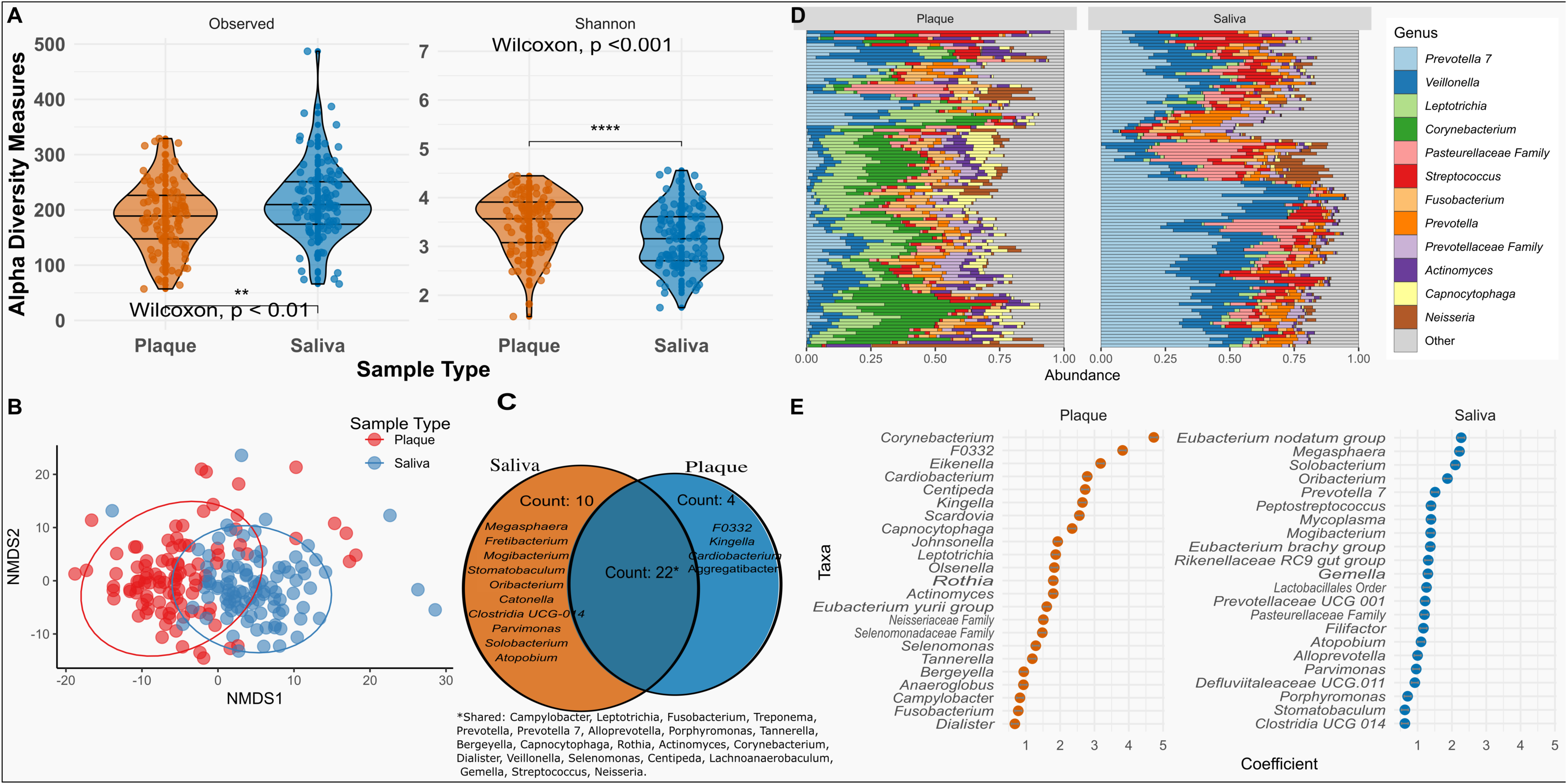
(A) Violin plots comparing alpha diversity measures (Observed richness and Shannon diversity) between plaque (orange) and saliva (blue) samples. Saliva samples exhibit significantly higher richness (Observed, Wilcoxon p < 0.01) and diversity (Shannon, Wilcoxon p <0.001) compared to plaque samples. (B) The NMDS plot illustrates that plaque samples (red points) form a distinct cluster, while saliva samples (blue points) are grouped separately. (C) Venn diagram showing the core microbiome and unique microbiome in saliva and plaque samples. (D) Relative abundance plot as stacked horizontal bars faceted by sample type (plaque and saliva). Genera category labelled “Other” was included to represent less abundant taxa. (E) Differential abundance plot showing taxa among saliva and plaque samples.

### The salivary and plaque oral microbiomes are uniquely shaped by the sociobiome

The plaque and saliva samples demonstrated significant associations with socioeconomic indicators (Table S2 and S3). Saliva samples uniquely showed significant difference in alpha diversity and beta diversity with age-related patterns (significant lowered in observed features as age increases, χ2: 6.13, p<0.04), secondary education level (Figure 2A-B, observed features: W=1730.5, p<0.01; Shannon index: W=1593, p<0.01), having Centrelink as income source (Figure 2C-D, observed features: W: 1478.5, p=0.02; Shannon index: W=1544, p<0.01), and HCC ownership (Figure 2E-F, observed features: W= 889, p=0.01; Shannon index: W=888, p=0.01). Active dental caries (Figure 2G-H) showed higher alpha diversity (observed: W=882.5, p=0.02; Shannon: W=766, p<0.01). In plaque samples, significant differences in both observed features and Shannon index were also found across age groups, particularly between younger (18-34 years) and older (>55 years) individuals (χ2: 6.08, p= 0.04 and W= 4.63, p=0.09). The observed features (W= 1627.5, p<0.01) and Shannon index (W= 1584, p = 0.01) were significantly higher for secondary education groups.

**Figure 2.**
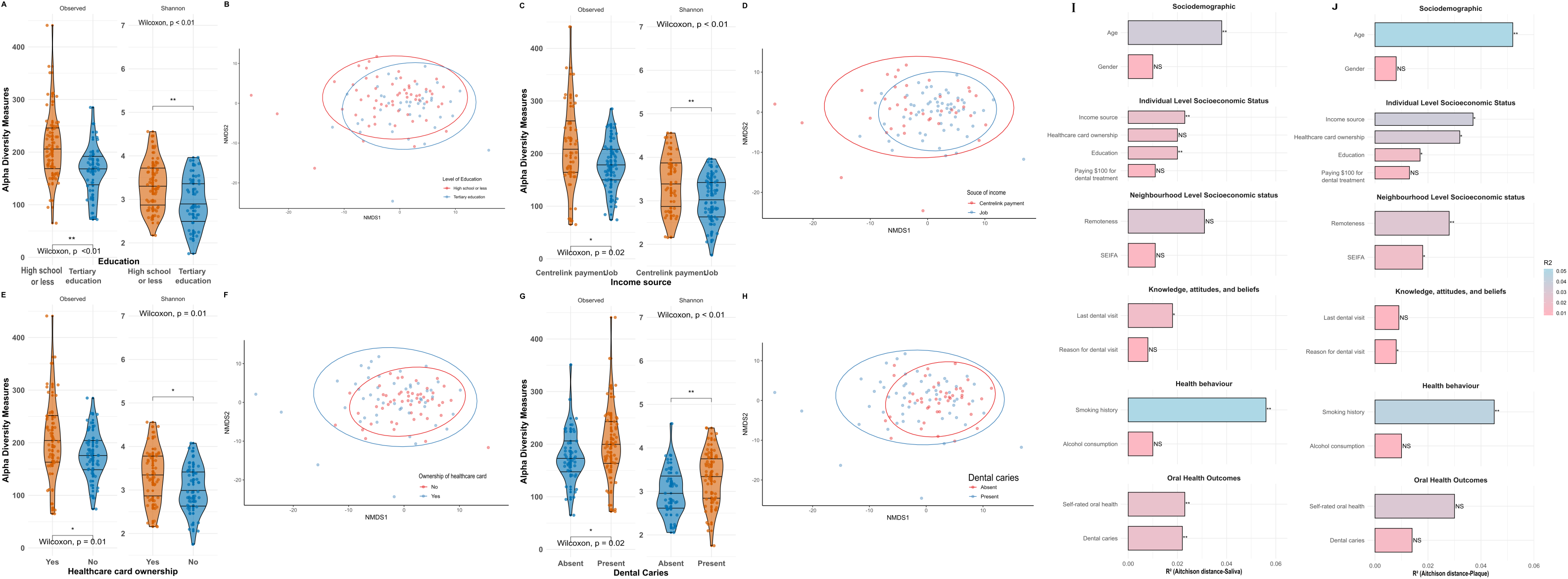
(A-B) Saliva samples for Violin plots showing alpha diversity measures (Observed features and Shannon index) of the salivary samples and NMDS plots based on Aitchison distances visualising beta diversity of the salivary microbiome by education level (< high school vs. tertiary), (C-D) income source (Centrelink payment vs. job), (E-F) healthcare card ownership (yes vs. no), and (G-H) dental caries status (present vs. absent). Each violin plot compares group distributions, with Wilcoxon p-values indicating statistical significance. For NMDS plots, the ellipses represent 95% confidence intervals for each group. (I-J) Bar plots summarising the proportion of variance (R²) in salivary (I) and plaque (J) microbiome composition explained by sociodemographic, individual and neighbourhood-level socioeconomic status, knowledge/attitudes, health behaviours, and oral health outcomes. Bars are coloured by R² value, with significant associations (p <0.05) highlighted.

The PERMANOVA using Aitchison distances revealed distinct microbial community structures in saliva (Figure 2I) and plaque samples (Figure 2J) associated with the sociobiome. In the saliva microbiome, the composition was significantly associated with age (F=1.94, R²=0.038, p<0.01), education (F=2.02, R²=0.020, p<0.01), income source (F=1.90, R²=0.038, p<0.01), last dental visit (F=1.79, R²=0.018, p=0.02), smoking history (F=2.85, R²=0.056, p<0.01), self-rated oral health (F=2.32, R²=0.023, p<0.01), and dental caries (F=2.28, R²=0.022, p<0.01). Similarly, PERMANOVA for plaque samples were significantly associated with age (F=2.69, R²=0.052, p<0.01), education (F=1.74, R²=0.017, p=0.04), income source (F=1.90, R²=0.037, p=0.01), remoteness (F=1.84, R²=0.018, p=0.02), smoking history (F=2.32, R²=0.045, p<0.01), and self-rated oral health (F=3.03, R²=0.03, p<0.01). Dental caries showed no association with plaque microbiome (p=0.09). These findings suggest that socioeconomic factors (education, income) and behavioural factors (smoking) consistently shape oral microbial communities across different oral habitats, while clinical manifestations like dental caries may have site-specific microbial signatures.

### Socioeconomic disadvantage and regional barriers drive dental caries risk

The RDA (Figure 3A) plot for saliva samples demonstrates that dental caries clusters closely with multiple markers of socioeconomic disadvantage, including Centrelink income, HCC ownership, regional residence, and low socioeconomic status. The clustering suggests these variables are strongly intercorrelated and collectively influence microbiome composition in similar directions. Problem-based dental visits align with the low SES cluster, reinforcing the connection between reactive (rather than preventive) dental care utilisation and socioeconomic disadvantage. Age-related patterns were evident, with older adults clustered downward. Overall, dental caries clustered with these socioeconomic disadvantage markers, and individuals with lower SES are more likely to seek dental care only when problems arise.

**Figure 3.**
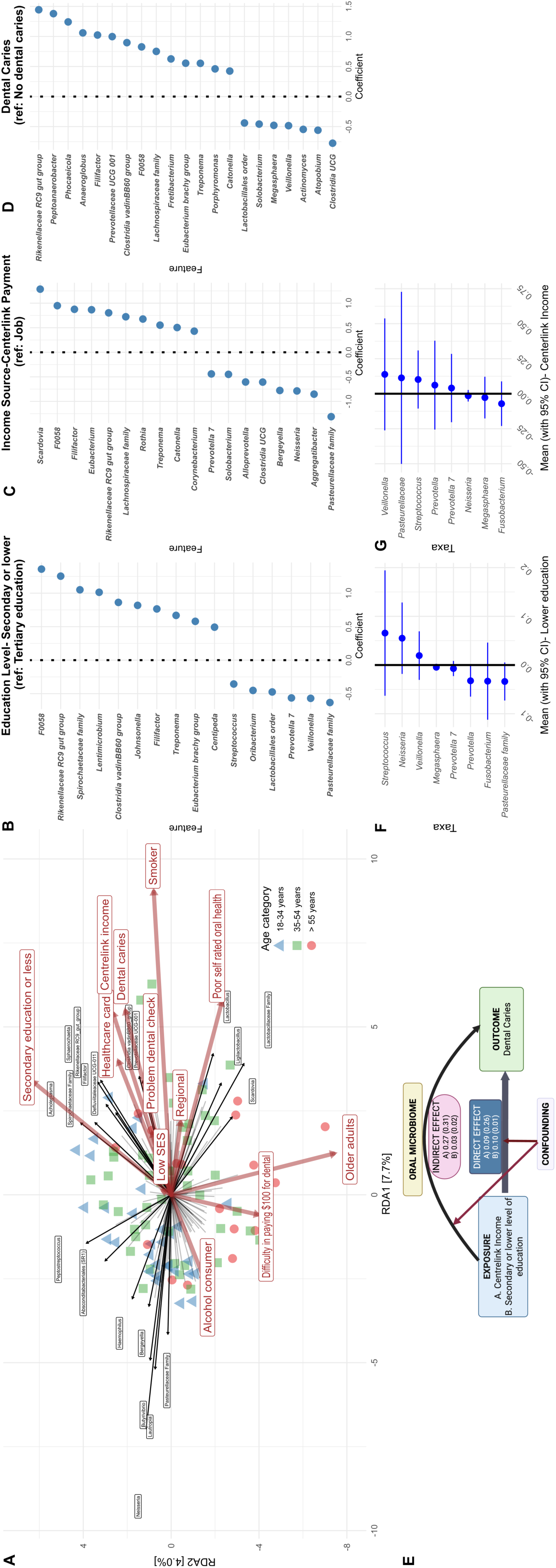
(A) Redundancy analysis (RDA) biplot of salivary microbiome composition showing associations with socioeconomic, behavioural, and clinical variables. Arrows represent explanatory variables, with direction and length indicating the strength and direction of association. Sample points are coloured by age category (green triangles: 18–34 years; blue squares: 35–54 years; red circles: >55 years). (B–D) Differential abundance analysis of salivary bacterial genera associated with (B) education level (secondary or lower vs. tertiary education), (C) income source (Centrelink payment vs. job), and (D) dental caries status (present vs. absent). Each plot displays the effect size (coefficient) for each taxon, with positive values indicating higher abundance and negative values indicating lower abundance relative to the reference group. (E) Causal mediation model illustrating the hypothesised pathway from socioeconomic exposure (Centrelink income or lower education) to dental caries outcome, mediated by oral microbiome composition. The model quantifies the direct effect (blue), the indirect microbiome-mediated effect (pink), and the role of confounding factors. (F–G) Forest plots showing the mean and 95% confidence intervals for the mediation effects of key bacterial taxa in the relationship between (F) lower education and dental caries, and (G) Centrelink income and dental caries. Positive values indicate taxa that mediate increased caries risk in disadvantaged groups; negative values indicate protective mediation.

A differential abundance analysis (Figure 3B-D, Table S4-S6) for saliva samples was plotted for education level, income source and dental caries. These forest plots illustrate how specific social determinants (education, income source) and clinical outcomes (dental caries) are significantly linked to variations in the abundance of multiple oral microbial taxa. The analysis revealed individuals with secondary education and those receiving Centrelink payments share several significantly abundant taxa that are also elevated in dental caries patients, particularly *Rikenellaceae RC9 gut group* (coefficients =1.05, 0.80, and 1.44 across education, income, and caries), *F0058* (coefficients = 1.36, 0.95, and 0.83), *Filifactor* (coefficients = 0.58, 0.88, and 1.02), and *Treponema* (coefficients = 1.01, 0.55, and 0.55). The findings support a mechanistic relationship between social factors, dental caries and the composition of the salivary microbiome.

### Lower income mediates microbiome composition, increasing dental caries risk

In mediation analysis, the direct effect (DE) of income on dental caries was 0.09 (SE=0.26), while the mediation effect (ME) operating through the oral saliva microbiome was 0.28 (SE=0.32). These results suggest that much of the impact of income on dental caries may be explained by changes in the oral microbiome (Figure 3 E-G, Table S7-S9). This suggests that approximately 75.6% (ME/TE = 0.28/0.37) of the income-caries relationship operated through changes in the oral microbiome composition. This suggests that lower income may lead to conditions that change oral bacterial communities in a way that promotes caries development. Similarly, mediation analysis was conducted for lower education and showed a significant direct effect (DE=0.10, SE=0.01) of lower education on dental caries risk independent of microbiome changes, and a smaller mediation effect (ME=0.03, SE=0.02) operating through microbiome alterations, yielding a total effect (TE=0.14, SE=0.03). These findings suggest roughly 21% (ME/TE=0.03/0.14) of this disparity was attributable to education-associated differences in oral microbiome composition, while 79% operated through direct pathways. Certain microbial taxa play varying roles in mediating the relationship between secondary or lower levels of education and dental caries. (Figure 3C, Table S7). *Streptococcus, Neisseria* and *Veillonella* demonstrated the strongest positive mediating effects, suggesting these genera significantly contributed to the pathway through which lower education leads to increased caries risk. Conversely, *Fusobacterium*, *Prevotella*, and *Pasteurellaceae* displayed a negative mediating effect (wide standard error suggests this estimate should be interpreted cautiously).

## DISCUSSION

Our study provides the first comprehensive analysis investigating how individual and neighbourhood socioeconomic factors shape the oral microbiome and mediate dental caries risk in an Indigenous Australian population. The findings reveal distinct microbiome profiles between plaque and saliva samples, with the salivary microbiome demonstrating stronger associations with socioeconomic indicators and dental caries status. This relationship suggests a potential biological pathway through which social determinants manifest as oral health inequities. Most notably, approximately 75% of the relationship between lower income (as indicated by Centrelink dependency) and dental caries risk is mediated through alterations in the oral microbiome. These findings have significant implications for understanding the biological mechanisms underlying persistent oral health disparities experienced by Indigenous communities, moving beyond behavioural explanations to recognise how structural disadvantage becomes biologically embedded through changes in the oral microbiome.

A key finding from our study is the differential impact of sampling methods on capturing socioeconomic associations. While plaque and saliva samples showed significant associations with age, education, income source, and smoking history, the saliva microbiome composition uniquely demonstrated significant relationships with healthcare card status, patterns of last dental visits, and the presence of dental caries. This indicates that saliva could be a better medium for exploring socioeconomic effects on oral microbiota and related health issues (17). This observation aligns with Handsley-Davis et al.’s (14) work with Indigenous children, where salivary characteristics were significantly associated with microbiota diversity and composition. The greater sensitivity of salivary samples to socioeconomic gradients may reflect saliva’s role as an integrative fluid representing the entire oral cavity rather than site-specific plaque biofilms, providing a more comprehensive view of the oral ecosystem shaped by various socioeconomic exposures over time (16, 17).

The RDA analysis further emphasised how socioeconomic disadvantage shapes salivary microbiome composition, with education levels emerging as the strongest determinant, followed closely by income source and healthcare card status. These socioeconomic indicators clustered with dental caries, suggesting potential mechanistic links between social disadvantage, microbial community structure, and oral disease outcomes. The differential abundance analysis provided further evidence for this pathway, identifying specific bacterial taxa (*Rikenellaceae RC9 gut group*, *F0058*, *Filifactor*, and *Treponema*) that were consistently associated in both low SES environments and individuals with dental caries. These patterns align with studies of Indigenous children in the Northern Peninsula Area (NPA) of Far North Queensland, where higher microbial diversity was linked to proxies for lower socioeconomic status and less frequent toothbrushing (14).

Mediation analysis revealed that approximately 75% of the relationship between income (Centrelink payment) and dental caries risk operates through changes in oral microbiome composition. Despite Centrelink cardholders having access to public dental care, high caries rates persist due to systemic gaps in preventive care, financial barriers, and underlying socioeconomic determinants. Public dental services often prioritise emergency treatments over preventive care, with waitlists exceeding 12–24 months in many regions. This gap between theoretical access and practical utilisation mirrors findings from previous Australian studies demonstrating that Indigenous Australians disproportionately experience poor oral health (19), with limited access to culturally appropriate and timely dental care being a significant factor (20). The microbiome-mediated effect was significantly smaller for educational attainment (21%), suggesting that different aspects of socioeconomic status may affect oral health through distinct causal pathways.

Our study has several methodological strengths. (1)Including both plaque and saliva samples enabled direct comparison of these sampling methods for detecting socioeconomic gradients. (2) The application of sophisticated mediation analysis allowed us to quantify the contribution of microbiome alterations to the overall relationship between SES and dental caries. (3) Focusing on Indigenous Australians addresses an important gap in the literature, as most oral microbiome research has been conducted in non-Indigenous populations despite evidence that Indigenous communities may have unique oral microbiota shaped by distinct genetic, cultural, and environmental factors (21, 22). However, there were a few limitations.

The sample size of 100 participants, while providing adequate power for detecting major patterns, may have limited our ability to identify more subtle associations or to comprehensively explore interactions between different dimensions of socioeconomic status. Additionally, while our mediation analysis suggests a substantial role for microbiome-mediated effects, the wide standard errors in these models indicate considerable uncertainty in the estimates.

The microbiome-mediated effects of income on dental caries risk suggest that interventions targeting microbial ecology, such as water fluoridation, professionally applied topical antimicrobials or prebiotics promoting beneficial bacteria, may help mitigate socioeconomic disparities in oral health outcomes (6). However, the concurrent finding that education’s impact operates primarily through non-microbiome pathways underscores the need for multipronged approaches addressing both biological and social determinants of oral health. Future research should build upon these findings through longitudinal studies tracking changes in oral microbiome composition, socioeconomic circumstances, and dental caries over time. Integrating metagenomic shotgun approaches would provide insights into functional changes in the microbiome beyond taxonomic shifts, potentially revealing specific metabolic pathways through which socioeconomic factors influence oral health (23). Our findings have important implications for addressing oral health disparities.

This study demonstrates that socioeconomic disadvantage significantly shapes oral microbiome composition in Indigenous Australians, with salivary microbiota serving as a sensitive indicator of these social gradients. The substantial contribution of microbiome-mediated effects to the relationship between income and dental caries highlights an important biological pathway through which social inequalities manifest as oral health disparities. These findings suggest that addressing Indigenous oral health inequalities requires both upstream approaches targeting social determinants and downstream interventions addressing the biological consequences of socioeconomic disadvantage on the oral microbiome.

## MATERIALS AND METHODS

This study is the baseline cross-sectional study of a larger ongoing 12-month follow-up study, with the primary goal of delivering culturally safe dental care to Indigenous South Australians. A detailed study protocol has been published (20). Ethical approval was obtained by the Indigenous Health Council of South Australia’s Human Research Ethics Committee (04-22-990) and the University of Adelaide Human Research Ethics Committee (20). The research followed the World Medical Association Declaration of Helsinki. This project adopted a decolonising methodology and collaborative community-based participatory action research approach, integrating Indigenous and Western research methods into a co-designed framework (24, 25). This approach emphasises Indigenous leadership, governance, and engagement in reflexive practice. This study was overseen by an Indigenous Oral Health Unit Reference Group (IOHURG), which provided advice on all aspects of project staff employment, participant recruitment, data collection, analysis and feedback (25). A written informed consent was obtained from the all the participants.

### Study design and data collection

Prior to data collection, all staff participated in cultural awareness sessions. The project was guided by an Aboriginal reference group. During engagement in Communities, the team followed the guidance of senior Aboriginal Elders on Community cultural protocols. Recruitment took place in metropolitan and regional areas of the Australian state of South Australia from July 2022 to December 2023, followed by a 12-month follow-up phase from March 2024 to March 2025. This study analysed the baseline data of 100 adults (that had microbial samples collected) from a convenience sample of 280 adult participants. Based on previous research (26, 27), recruitment methods included self-nomination, word of mouth, flyers and posters, and personal visits to local community centres and health clinics. Eligibility criteria consisted of participants being Indigenous or Torres Strait Islander or both, over 18 years of age, and living in South Australia. All participants received written and verbal information about the study, and written informed consent was obtained.

The data collection and dental examination were conducted in participants’ homes, workplaces, community centres, or dental clinics, adapting to each participant’s most convenient and comfortable setting. Data were collected by experienced Indigenous and non-Indigenous research officers, all of whom were trained and calibrated to deliver the study’s aims, self-reported questionnaires, and point-of-care (POC) testing for general health checks (20). The research team prioritised cultural sensitivity throughout the study design and implementation. Indigenous research officers administered self-reported questionnaires, which were completed as interviews, on paper, or online via REDCap (Research Electronic Data Capture software (28).

### Dental Examination

Two calibrated dental practitioners (SN and KK) performed comprehensive oral assessments using head-LED lights, portable dental equipment, and single-use dental instruments. The examination included a five-surface evaluation of each tooth, employing a coding system to record as either sound teeth (S) or decay using ICDAS II scores (D). An ICDAS II score of 4 or higher was categorised as having caries present (29).

### Variables

The categorisation of each variable is described in detail in Supplementary Text S1 and consists of five domains. The sociodemographic domain included age, sex, and residential location. The SES domain included both individual-level and neighbourhood-level indicators. Four measures were considered for individual-level SES: education level (secondary vs tertiary), income source (job vs. Centrelink), ownership of a health care card (HCC) and difficulty paying $100 for dental treatment (hard vs not hard). Neighbourhood-level SES was assessed using the Socio-Economic Indexes for Areas (SEIFA) and remoteness score (30, 31). The knowledge, attitude and beliefs domain included time since the last dental visit (<1 year or >1 year), the reason for visiting the dentist (check-up or problem-based), and cost-related avoidance of dental visits (yes or no). The health behaviour factors included self-reported diabetes smoking habits (current, past, or non-smoker) and alcohol consumption and frequency (never, monthly, or weekly). Oral health outcomes factors included self-rated oral health (good or poor) and dental caries status (present or absent).

### Sample collection

Saliva and plaque samples were collected by trained and calibrated dental practitioners (SN & KK). Unstimulated 2 ml saliva samples were collected using a saliva collection tube (Zymo DNA/RNA Shield Saliva Collection Kit, Zymo Research, California, USA). The supragingival plaque was collected from six pooled sites using a sterile single-use universal Gracey curette following a standardised protocol (32) and carefully transferred into a 2 mL tube containing a DNA/RNA shield (Zymo Research, California, USA). Additional control samples, such as environmental (ENV) and curette wash (CW), were collected. All the samples were transported into the laboratory for storage at –20°C and then shipped to the microARCH Lab at Pennsylvania State University for 16S rRNA amplicon sequencing; this facility is designed to reduce contaminant signatures using specific workflows, UV radiation, and positive pressure. All work was conducted in this laboratory in a still air hood. The entire protocol for biological and control sample collection is described in Supplementary Text S2.

### Microbial analysis and data pre-processing

DNA was extracted from all the biological samples using the MagMax Microbiome Ultra Nucleic Acid Isolation Kit (Applied Biosystems). Extraction blank controls (EBCs) and no template controls (NTCs) were also processed alongside to identify background laboratory contamination. The V4 region of the 16S rRNA gene was amplified, purified, and sequenced on an Illumina MiSeq platform, as previously described (33). This method has been previously used to amplify Indigenous Australian oral microbiomes using the 16S rRNA V4 region, with special attention paid to contaminant control, inclusion of contaminant monitoring, and clean-room protocols (34). Supplementary Text S3 describes the complete DNA extraction, amplification, and sequencing process, as previously adapted from the Caporoso method (35). Data pre-processing was performed in QIIME2 software (v.2024.11) (36), which involved demultiplexing, quality filtering, and denoising reads to generate amplicon sequence variants (ASVs) (33). Sequences were aligned with MAFFT to create phylogenetic trees and a rooted phylogenetic tree. The Silva 138 (99% of full-length sequences) database was used for taxonomic assignment to ASVs. The detailed protocol for bioinformatics analysis is outlined in Supplementary Text S4.

Contaminant removal was performed in two sequential steps for comprehensive contaminant detection and removal. First, we compared EBCs (n=40) and NTCs (n=6) samples against the biological samples (Figure S1, S2) and identified 54 taxa that were more abundant in controls and biological samples (Table S10); all of these taxa were removed from the data set. Next, we compared ASVs within the ENV (n=20) and CW (n=2) controls using *decontam* (37) (Figure S2) and detected an additional 39 contaminants (Table S11); these taxa were also removed from the data set. After both decontamination steps, microbiota counts, metadata, and taxonomy artefacts were imported into RStudio to create a *phyloseq* object for further analysis. Data filtering was performed in multiple stages to refine the microbiome dataset. Initially, ASVs with zero counts across all samples were removed. Subsequently, rare taxa were filtered out based on prevalence (<5% of samples) or total abundance (<10 reads), along with known contaminant sequences identified as chloroplast or mitochondria. Saliva samples had sequencing depths ranging from 17,692 to 95,253 reads, with a median of 56,456 and a mean of 55,005. Plaque samples exhibited sequencing depths ranging from 12,362 to 88,115 reads, with a median of 62,042 and a mean of 60,140. After filtering, saliva samples contained 678 and 544 ASVs.

## Statistical analysis

### Comparison of saliva and plaque samples

All the analysis was conducted in R version 4.4.2 (2024-10-31), and the script can be found at https://github.com/sonianath/Indigenous_Australian_Oral_Microbiome. Alpha diversity measures of the microbiome were analysed for two sample types: plaque and saliva, after the data were rarefied to even sampling depth, where the rarefaction curve reached a plateau (Figure S3). The metrics used were “Observed” (observed species, the number of observed taxa) and Shannon’s diversity (a measure of microbial diversity incorporating both richness and evenness). Violin plots were generated to visualise the distribution of alpha diversity for each sample type and compared using the Wilcoxon rank-sum test. Beta diversity was visualised using non-metric multidimensional scaling (NMDS) using the *MicroViz* package (38), using Aitchison’s distances. A compositional bar plot was generated to visualise the relative abundance of microbial genera in plaque and saliva samples aggregated at the genus level, retaining the top 12 most abundant genera. Saliva and plaque samples were compared for core microbiome genera, defined as those present in at least 80% of samples with a minimum abundance of 5 counts. The core genera unique to saliva and plaque and shared between both sample types were determined, and a Venn diagram was created (39). Lastly, differentially abundant bacterial genera were identified in saliva and plaque samples, controlling for age and sex using the negative binomial model for count data and cumulative sum scaling (CSS) normalisation from the *MaAsLin2* package (40).

### Analysing the impact of socioeconomic factors on oral microbiome from the plaque and saliva samples

Redundancy analysis (RDA) assessed the relationships between microbial community composition and explanatory variables using the *MicroViz* package (38). Explanatory variables included demographic factors (e.g., age categories, female gender), socioeconomic indicators (e.g., low SES, secondary education or less, Centrelink/social welfare as an income source, healthcare card ownership, having difficulty paying $100 for dental treatment), behavioural factors (e.g., smoking, alcohol consumption) and oral health factors (poor self-rated oral health, dental caries. The ordination plot was generated to visualise the clustering of samples and the directional influence of variables on microbial community composition grouped by age categories (18–34 years & 35–54 years), with taxa and explanatory variables represented as vectors, with their length and direction indicating their strength and influence (41). Similarly, the alpha diversity (Observed and Shannon’s) and beta diversity (Aitchison’s) were compared separately across independent variables for plaque and saliva samples after rarefaction to even depth. For alpha diversity, either the Wilcoxon test for two groups or the Kruskal-Wallis test for multiple groups was used, followed by the post hoc Dunn test for pairwise comparison controlling for false discovery rates using the Benjamini-Hochberg method (p<0.05). For beta diversity, NMDS plots were used for comparison, and pairwise PERMANOVA was used to test the difference between the groups. A multivariate PERMANOVA included sociodemographic, SES, knowledge, attitude indicators, and health behaviour variables, and health behaviour variables were compared to explain the variance of each variable. Only the SES variables that were significant in beta diversity were analysed for differential abundance, along with dental caries (only for saliva samples), controlling for age and sex (40).

### Mediation analysis

Mediation analysis was used to examine how income source (Centrelink vs Job) and education level (high school or less vs. tertiary) affect dental caries, with the oral microbiome as a potential mediator. Income source and education were selected as primary exposures, and dental caries as the outcome due to their significant impact observed in the PERMANOVA (adonis) analysis of saliva samples. First, propensity score matching was conducted as a preprocessing step to balance confounding variables between educational groups, implementing nearest neighbour matching with replacement (1:8 ratio) based on age, sex, income, healthcare card status, smoking, difficulty paying $100 for dental treatment, dental visit frequency, remoteness (location), and alcohol consumption (42). This matching process created comparable groups for analysing the causal pathways from education to dental caries using the *MatchIt* package (43). We evaluated the balance of covariates across the groups using summary statistics and visualised through a love plot (Figure S4), showing standardised mean differences before and after matching. We then applied the *SparseMCMM* package (44) with the matched data, using level of education as the treatment variable, dental caries presence/absence as the outcome, and oral microbiome composition (filtered at a genus level using 0.1% abundance and 5% prevalence thresholds) as the mediator. A 100 random data splits were incorporated to enhance robustness, and this was used to calculate the direct effects, indirect effects, and total effects. A component-wise effect of each taxon was calculated as a mediator.

## ETHICAL APPROVAL AND CONSENT TO PARTICIPATE

Ethical approval was obtained by the Indigenous Health Council of South Australia’s Human Research Ethics Committee (04-22-990) and the University of Adelaide Human Research Ethics Committee. The research followed the World Medical Association Declaration of Helsinki. This study was overseen by an Indigenous Oral Health Unit Reference Group (IOHURG), which provided advice on all aspects of project staff employment, participant recruitment, data collection, analysis and feedback. A written informed consent was provided by all the study participants.

## CONSENT FOR PUBLICATION

All data included in this manuscript have been approved for publication, and no additional consent is required.

## DATA AVAILABILITY STATEMENT

All sequence data generated in this project are available at NCBI under BioProject ID PRJNA1303862. The R scripts to analyse the 16S rRNA amplicon sequencing data are available at https://github.com/sonianath/Indigenous_Australian_Oral_Microbiome.

## COMPETING INTEREST

The authors declare that they have no known competing financial interests or personal relationships that could have appeared to influence the work reported in this paper.

## FUNDING

This research was supported by the National Health and Medical Research Council Research Fellowship (APP1102587).

## AUTHOR CONTRIBUTIONS

S.N: contributed to conception and design, contributed to the acquisition, drafted manuscript, and critically revised the manuscript and gave final approval. L.W: contributed to design, contributed to interpretation, drafted manuscript and critically revised manuscript. G.G: contributed to conception and design, contributed to interpretation, drafted manuscript, and critically revised the manuscript and gave final approval. J.H: contributed to design, contributed to analysis and interpretation, drafted manuscript, and critically revised the manuscript and gave final approval. M.T: contributed to conception and design, contributed to interpretation, drafted manuscript, and critically revised the manuscript and gave final approval. K.K: contributed to conception and design, contributed to interpretation, drafted manuscript, and critically revised the manuscript and gave final approval. LJ: contributed to conception and design, contributed to funding acquisition, analysis, and interpretation, drafted manuscript, and critically revised manuscript. All authors gave their final approval and agreed to be accountable for all aspects of the work.

## ACKNOWLEDGEMENTS

We respectfully acknowledge the Kaurna people, the Ngarrindjeri people, the Adnyamathanha people, the Nukunu people, the Barngarla people, and all other First Nations of those involved in this project. We gratefully acknowledge the participants, their families, and communities for their invaluable contributions to this project. We thank the IOHU team and students for their dedicated assistance with fieldwork. Our appreciation extends to Sonder, Closing the Gap, Tiraapendi Wodli, Neporendi, Taikurrendi Childcare, Christies Beach High School, Nuriootpa High School, and Kanggawodli for their support and collaboration. Special thanks to Pedro for assisting with statistical analysis and to Nicole Moore for supporting the laboratory analysis and management.

